# p63 and PITX1 sustain a pre-invasive malignant keratinocyte population in squamous cell carcinoma precursors

**DOI:** 10.64898/2026.05.21.725073

**Authors:** Ramon Staeger, Aizhan Tastanova, Adhideb Ghosh, Paul Gueguen, Isabel Kolm, Egle Ramelyte, Evelyn Lattmann, Veronika Haunerdinger, Tugay Karakaya, Marta Slaufova, Michela Di Filippo, Hans-Dietmar Beer, Mitchell P. Levesque, Reinhard Dummer

**Affiliations:** Department of Dermatology, University Hospital Zurich, Zurich, Switzerland; Medical Faculty, University of Zurich, Zurich, Switzerland; Functional Genomics Center Zurich, University of Zurich and ETH Zurich, 8057 Zurich, Switzerland; Department of Pathology, Luzerner Kantonsspital, Lucerne, Switzerland

## Abstract

**Background:** Cutaneous squamous cell carcinoma (cSCC) is among the most common human cancers, yet the cellular identity and molecular programs of its preinvasive precursor, actinic keratosis (AK), remains poorly defined.

**Methods:** We applied CITE-seq to patient-matched AK, UV-exposed normal skin, and non-UV-exposed normal skin (n=5 patients, 12 biopsies) and performed spatial whole-transcriptome profiling in an independent cohort (n=4) to map pre-invasive keratinocyte states at single-cell resolution.

**Results:** We identify AK-specific keratinocytes (ASK), a discrete population localized to the dysplastic basal epidermis and characterized by UV-associated mutational signatures (SBS7b), high mutational burden, and recurrent copy number alterations including 9p loss and 8q gain. ASK occupies a basal-like undifferentiated state sustained by a ΔNp63/PITX1 regulatory module that attenuates Notch/HES1-driven differentiation and activates glycolytic metabolism. Comparison with published cSCC data reveals that ASK share core tumor-propagating gene networks with tumor-specific keratinocytes (TSK), including IGFBP6, IGFBP2, and ITGA6, but lack invasion effectors MMP1, MMP10, and PTHLH. Functional experiments identify IGFBP6 as a pro-proliferative factor in AK-derived keratinocytes. The AK microenvironment shows expansion of inflammatory basal keratinocytes, barrier disruption, and early immunosuppressive T cell remodeling.

**Conclusions:** These findings define the molecular identity of a pre-invasive malignant keratinocyte population governed by p63/PITX1 and distinguish early oncogenic programs shared with invasive cSCC from later-acquired invasion effectors, identifying candidate targets for prevention or treatment of squamous cell carcinoma.

## Introduction

Cutaneous squamous cell carcinoma (cSCC) is among the most common human malignancies, arising on a background of chronic ultraviolet (UV) exposure and field cancerization (Leiter et al., 2023; Ratushny et al., 2012). Actinic keratosis (AK) represents an early intraepidermal stage of squamous neoplasia, characterized by clonal expansion of atypical, frequently p53 mutated keratinocytes confined to the epidermis (Criscione et al., 2009; Czarnecki et al., 2002; Hofbauer et al., 2014; Werner et al., 2013). Epidemiological and histopathological data consistently support AK as the dominant precursor route to invasive cSCC (Marks et al., 1988; Mittelbronn et al., 1998), making it a uniquely accessible *in vivo* model for studying early squamous carcinogenesis.

UV-driven mutagenesis is central to this process. Sequencing studies have revealed recurrent alterations converging on impaired differentiation and deregulated cell-cycle control, including inactivating events in *TP53*, *CDKN2A*, *NOTCH1/2*, and activating mutations in RAS pathway genes (Inman et al., 2018; Pickering et al., 2014; South et al., 2014; Thomson et al., 2021). UV exposure additionally remodels the stromal and immune microenvironment through chronic inflammation and extracellular matrix reorganization (Bachelor & Bowden, 2004; Dotto, 2014; Goruppi & Dotto, 2013; Timares et al., 2008). How these genetic and microenvironmental inputs reorganise epithelial cell states at the pre-invasive stage remains incompletely understood. Specifically, it is unclear which keratinocyte programs mark malignant commitment and which features distinguish early malignant keratinocytes from invasive tumor states.

Single-cell RNA sequencing (scRNA-seq) studies of invasive cSCC have identified a tumour-specific keratinocyte (TSK) population associated with extracellular matrix remodelling and invasion, and implicated p63-centred transcriptional networks and glycolytic metabolism as core dependencies of squamous tumour growth (A. L. Ji et al., 2020). Two subsequent single-cell analyses of AK reported disease-associated keratinocyte states, upregulation of markers such as *CDKN2A* and *IGFBP2*, and roles for epithelial–stromal crosstalk; chromatin accessibility profiling further implicated p63-mediated transcriptional activation as a driver of squamous progression (J. Li et al., 2023; D.-D. Zou et al., 2023). However, a patient-matched, multimodal characterization of AK with spatial resolution has not been reported.

Here, we apply simultaneous single-cell RNA and surface proteome sequencing (Lischetti et al., 2023; Stoeckius et al., 2017) to matched AK, UV-exposed normal skin, and non-UV-exposed normal skin from five patients. We then integrate these data with spatial whole-transcriptome profiling in an independent cohort to anchor disease-associated cell states within tissue architecture. This framework identifies a transcriptionally and genomically defined population of malignant keratinocytes in AK characterized by malignant genomic hallmarks, a basal-like undifferentiated phenotype, and selective ΔNp63 and PITX1 regulatory activity. By defining the molecular identity and regulatory circuitry of this early malignant state, and relating it to the invasive TSK program in cSCC, this study sheds light on the cellular events underlying early squamous carcinogenesis.

## Results

To delineate cellular programs in actinic keratosis (AK), we profiled 12 skin biopsies from five patients with single cell whole transcriptome sequencing coupled to simultaneous quantification of 130 surface proteins using CITE-seq (Cellular Indexing of Transcriptomes and Epitopes by Sequencing) (**Figure 1A**) (Stoeckius et al., 2017). The cohort comprised matched pairs of AK (n=4), UV-exposed normal skin (UVES, n=5), and non-UV-exposed normal skin (NUVES, n=3). Viable cryopreserved tissues were dissociated to single cells under a uniform workflow (Burja et al., 2022). To anchor these cell states in tissue architecture, we also generated spatial whole transcriptome maps from an independent patient cohort (n=4) (**Figure 1A**).

**Figure 1.**
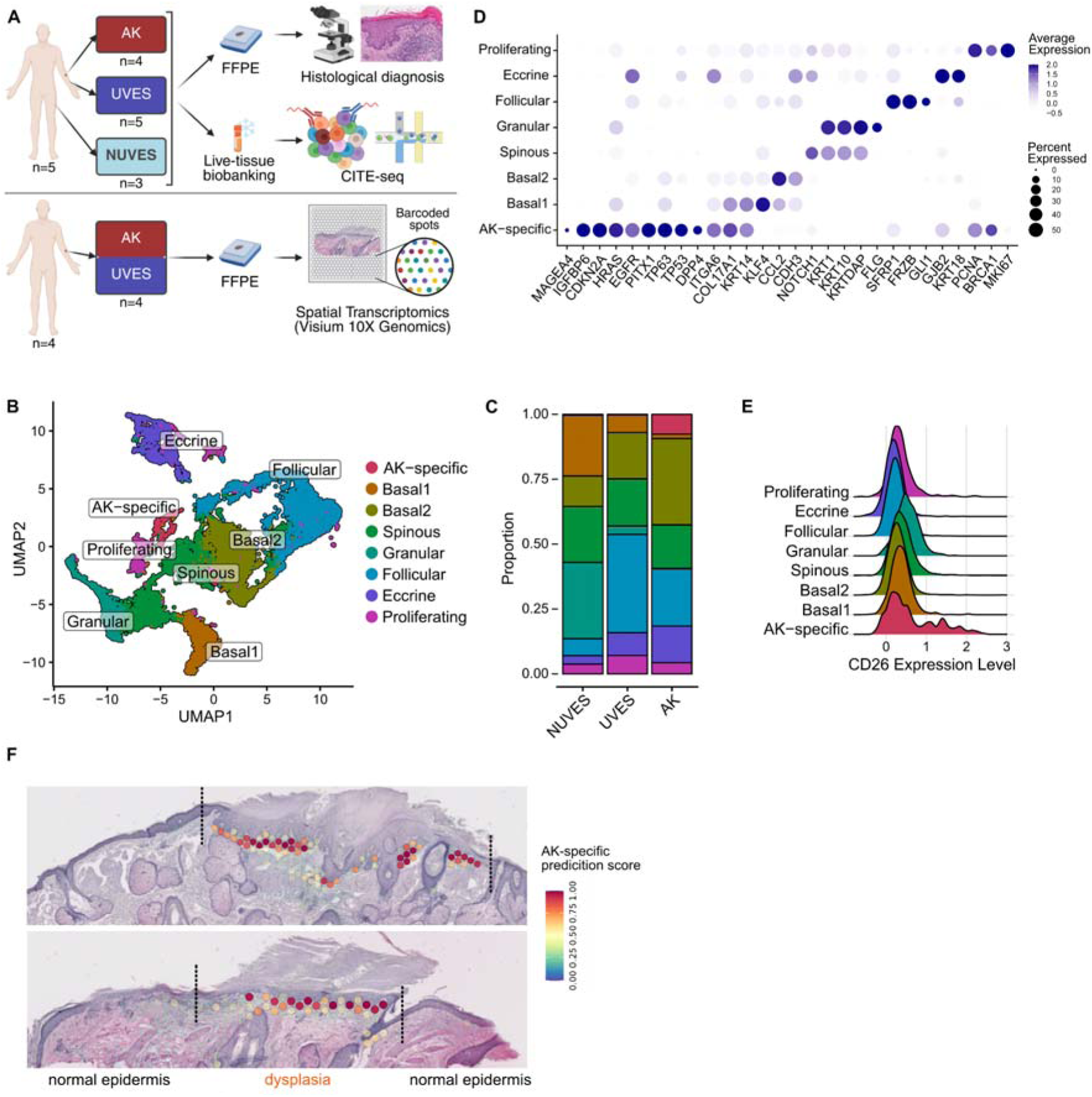
**(A)** Study design. **(B)** UMAP plot of integrated keratinocyte subclusters. **(C)** Proportion of keratinocyte subclusters by condition. **(D)** Dot plot of marker gene expression by subcluster. Dot size reflects the fraction of expressing cells; color reflects average expression. **(E)** CD26 surface protein expression by subcluster shown as ridge plot. **(F)** Spatial deconvolution of ASK in AK tissue sections. Dysplastic and morphologically normal epidermis are indicated. AK: Actinic keratosis. ASK: AK-specific keratinocytes. UVES: UV-exposed normal skin. NUVES: Non-UV-exposed normal skin. FFPE: Formalin-fixed paraffin-embedded. CITE-seq: Cellular Indexing of Transcriptomes and Epitopes by Sequencing. UMAP: Uniform Manifold Approximation and Projection.

### 1. Integrated single-cell and spatial profiling identifies an AK-specific keratinocyte population

Unsupervised clustering of keratinocytes based on single-cell transcriptomic data revealed eight distinct subpopulations (**Figure 1B**). Among these, one subcluster was highly enriched in AK lesions, comprising 7.6% of keratinocytes compared to only 0.2% and 0.4% in UVES and NUVES control skin, respectively (**Figure 1C**). This population was termed AK-specific keratinocytes (ASK). ASK were characterized by high expression of oncogenic and regulatory genes, including *MAGEA4* (a cancer-testis antigen) (Ishihara et al., 2020), *IGFBP6*, *CDKN2A*, *HRAS*, and *EGFR* (**Figure 1D**). Notably, the transcription factors *PITX1* and *TP63* were significantly upregulated in this population (**Figure 1D**). ASK were further marked by significant *DPP4* gene expression (p-adj<10^-76^, log2FC=3.77), confirmed at the protein level by surface overexpression of CD26 (p-adj<10^-9^, log2FC=1.65) (**Figure 1D, E**).

To determine the spatial context of ASK cells, we performed spatial whole transcriptome analysis on an independent AK cohort (n=4). Deconvolution using the scRNA-seq derived cell typing mapped ASK cells to the basal layer of the dysplastic epidermis, while morphologically normal epidermis lacked the signature (**Figure 1F**). Thus, integrated single-cell and spatial analyses establish ASK as a discrete keratinocyte population uniquely associated with the dysplastic compartment of AK.

To validate this finding, we projected ASK onto published scRNA-seq datasets of AK and cSCC (Hao et al., 2021; A. L. Ji et al., 2020; D.-D. Zou et al., 2023). Cells matching the ASK phenotype (prediction score ≥95th percentile) were identified in both datasets, where they progressively increased from normal skin to premalignant (p<0.02) and malignant (p<0.03) stages (**Supplementary Figure 1A and B**). This confirms the involvement of ASK cells throughout squamous carcinogenesis.

The remaining keratinocytes segregated into three broad categories based on differentiation status and anatomical associations. The first group comprised less differentiated cells including two basal clusters (basal1 and basal2) characterized by the expression of *COL17A1, KRT14,* and *CCL2*; and a proliferating cluster (*PCNA*, *MKI67*). The second group consisted of differentiating keratinocytes, represented by a spinous cluster (*KRT1, KRT10, NOTCH1*), and a granular cluster (*KRTDAP, FLG*). The third category included keratinocytes of skin appendages, including follicular keratinocytes (*SFRP1, FRZB, GLI1*) and eccrine gland cells (*GJB2, KRT18*) (**Figure 1B-D**).

### 2. ASK harbor malignant genomic hallmarks

Genomic characterization using inferred large-scale copy number variation (iCNV) confirmed typical copy number alterations (CNA) previously identified in AK lesions to be specifically present in ASK cells (Kim, Jung, et al., 2023; Thomson et al., 2021). These CNAs included loss of chromosome 9p (harboring the tumor suppressor *CDKN2A*) and gain of chromosome 8q (harboring the oncogene *MYC*) (**Figure 2A-C**). These findings suggest that ASK represent a malignant cell population within AK lesions.

**Figure 2.**
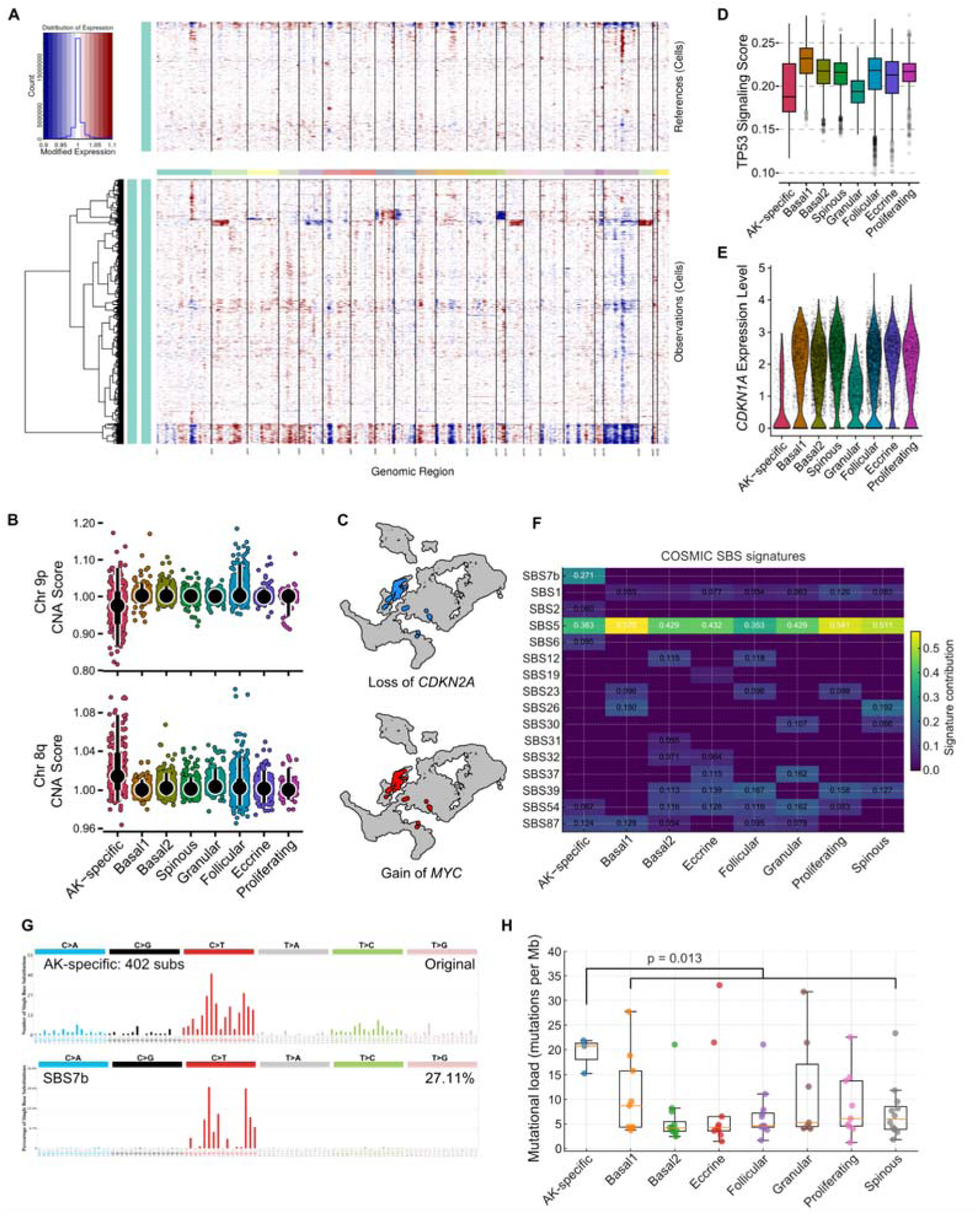
**(A)** inferCNV heatmap of keratinocytes from AK. Genomic regions are shown along the x-axis; individual cells along the y-axis. **(B)** CNA scores for chromosome 9p and 8q by keratinocyte subcluster. **(C)** UMAP highlighting cells with inferCNV-inferred loss of *CDKN2A* (chr9p) or gain of *MYC* (chr8q). **(D)** TP53 Hallmark signaling scores by subcluster (Liberzon et al., 2015). ASK scored significantly lower than all subclusters (Wilcoxon rank-sum test, p-adj<10⁻) except granular (p-adj=0.78). **(E)** *CDKN1A* expression by subcluster. **(F)** COSMIC single-base substitution (SBS) signature contributions across keratinocyte subclusters shown as heatmap. Values reflect the fraction of mutations attributed to each signature; values ≥0.05 are shown. **(G)** SBS signature reconstruction for the ASK mutational profile, showing a 27.1% contribution of the UV-associated SBS7b signature. **(H)** Mutational load (mutations per megabase) by subcluster, derived from SComatic calls on scRNA-seq data. ASK show significantly higher mutational load than other subclusters pooled (Mann-Whitney U test, U=185.0, p=0.013). Pairwise tests further supported higher load versus Basal2, Follicular, Spinous, and Proliferating (Benjamini–Hochberg, FDR ≤ 0.046), with non-significant trends versus Basal1, Eccrine, and Granular (FDR ≤ 0.105). AK: Actinic keratosis. ASK: AK-specific keratinocytes. CNA: Copy number alterations. CNV: copy number variations. UMAP: Uniform Manifold Approximation and Projection.

Additionally, ASK exhibited significant *TP53* overexpression (p-adj<10^-6^, log2FC=0.88) (**Figure 1D**), although pathway analysis indicated reduced p53 signaling compared to other keratinocyte clusters, except granular (p-adj<0.05) (**Figure 2D**) (Liberzon et al., 2015). In line with diminished p53 signaling, *CDKN1A* (p21), a key cell cycle regulator downstream of p53 (el-Deiry et al., 1993), was strongly downregulated in ASK (p-adj<10^-64^, log2FC=-2.03) (**Figure 2E**). This paradoxical pattern is consistent with loss-of-function mutations in the p53 DNA-binding domain, which often result in p53 overexpression, a phenomenon commonly observed in AK (Blokx et al., 2003; Nelson et al., 1994; Sung et al., 2022).

Somatic variants were called directly from scRNA-seq using SComatic (Muyas et al., 2024), then decomposed into COSMIC mutational signatures (Tate et al., 2019). In ASK, the UV-associated single-base substitution (SBS) signature 7b was detected with a contribution of 27.1%, with high reconstruction concordance (cosine similarity 0.973) **(Figure 2F-G, Supplementary Figure 2A)** (Hayward et al., 2017). Somatic variants in other keratinocyte subsets were predominantly explained by the background SBS5 signature, consistent with observations in non-neoplastic tissues **(Figure 2F)** (Moore et al., 2021). These mutational imprints identify ASK as the UV-damaged epithelial population within AK, consistent with genomic features characteristic of squamous carcinogenesis.

To directly compare genomic damage across keratinocyte populations, somatic variants were called from scRNA-seq with SComatic and summarized as mutational load (mutations per megabase, muts/Mb) per subcluster (Muyas et al., 2024). ASK displayed the highest mutational load (median 20.75 muts/Mb, range 15.26-21.84), whereas other keratinocyte populations had significantly lower mutational loads (medians 4–9 muts/Mb; p=0.013) (**Figure 2H**). For comparison, bulk sequencing studies reported a median of 10.4 muts/Mb in UVES (Kim, Bang, et al., 2023), 22.3 muts/Mb in AK lesions from immunocompetent patients (Thomson et al., 2021), and 50-60 muts/Mb in invasive cSCC (Pickering et al., 2014; South et al., 2014), placing ASK within the expected trajectory from premalignant to malignant keratinocytes.

### 3. ASK display a basal-like undifferentiated state and are regulated by **Δ**Np63 and PITX1

Integrated transcriptomic and surface-proteomic analyses positioned ASK within a basal, undifferentiated keratinocyte compartment. Surface proteomics showed elevated CD49f (integrin α6; encoded by *ITGA6*), a canonical basal keratinocyte marker, on ASK, basal1 and basal2 (p-adj<0.001, log2FC>0.30) (A. Li et al., 1998; Terunuma et al., 2007), with concordant *ITGA6* mRNA upregulation (**Figure 3A and 1D**). To position ASK within the keratinocyte continuum, transcriptomic similarity between subclusters was quantified using pairwise mean-squared-error (MSE) distances on scaled single-cell expression profiles, where lower MSE indicates greater similarity (**Figure 3B**) (Y. Ji et al., 2023). ASK most closely resembled proliferating and basal keratinocytes and were furthest from spinous/granular states; the ordering remained robust in Monte Carlo subsampling (100 iterations, 80% of cells per iteration; **Supplementary Figure 3A**). Marker-based scoring further corroborated enrichment of undifferentiated programs (keratins 5 and 14, cadherin 3) depletion of differentiation markers (keratin 1 and10, cadherin 1, desmoglein 1) in ASK (**Figure 3C**) (Wang et al., 2020). Consistently, integration of surface-protein and RNA data using weighted nearest neighbors (WNN) positioned ASK adjacent to basal/proliferative keratinocytes, confirming that this basal-like proximity holds across modalities (**Figure 3D**). Consistent with this undifferentiated transcriptional state, the AK-derived PM1 cell line, which harbors mutations in the known cSCC driver genes *TP53*, *NOTCH1/2*, and *CDKN2A (Hassan et al., 2019)*, failed to upregulate differentiation markers keratin 10, filaggrin, and loricrin under *in vitro* differentiation conditions (**Supplementary Figure 3B**).

**Figure 3.**
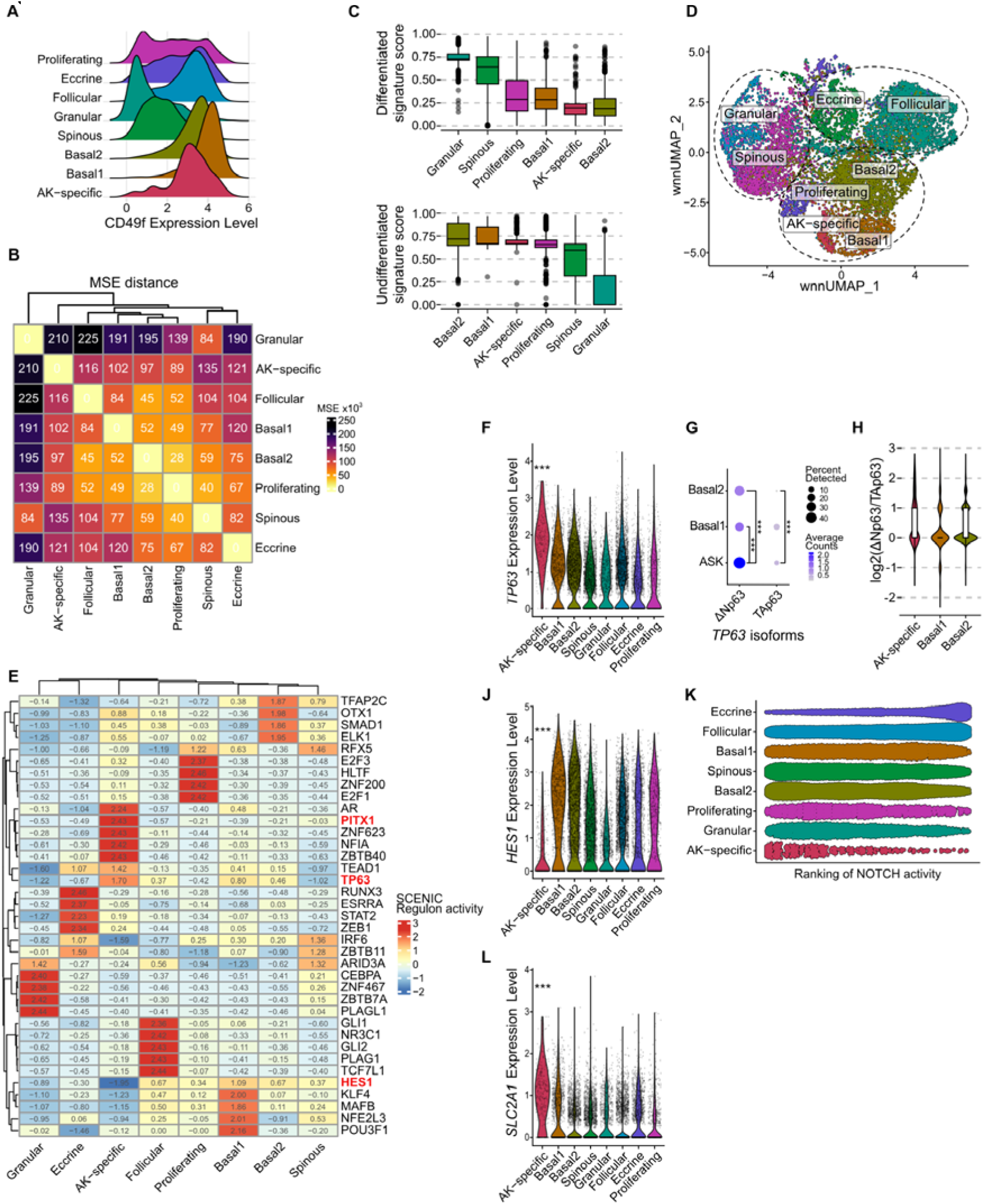
**(A)** CD49f surface protein expression by subcluster shown as ridge plot. **(B)** Pairwise mean squared error (MSE) distance heatmap between keratinocyte subclusters; lower values indicate greater transcriptomic similarity. **(C)** Enrichment scores of differentiated (*KRT1*, *KRT10*, *DSG1*, *CDH1*) and undifferentiated (*KRT5*, *KRT14*, *CDH3*) keratinocyte signatures by subcluster (Wang et al., 2020). **(D)** Weighted nearest neighbor (WNN) UMAP integrating RNA and surface protein information. **(E)** Scaled SCENIC regulon activity across keratinocyte subclusters. PITX1, TP63, and HES1 are highlighted. **(F)** *TP63* expression by subcluster. **(G)** ΔNp63 and TAp63 isoform expression in ASK and basal subclusters (Wilcoxon rank-sum test). **(H)** Log2 ratio of ΔNp63 to TAp63 isoform counts by subcluster. **(J)** *HES1* expression by subcluster. **(K)** Per-cell ranking of NOTCH Hallmark signaling enrichment scores by subcluster shown as beeswarm plot (Liberzon et al., 2015); ASK ranked significantly lowest (Wilcoxon rank-sum test, p-adj<10⁻¹). **(L)** *SLC2A1* (GLUT1) expression by subcluster. *\*\*\*: p<0.001, ASK: AK-specific keratinocytes*.

To determine which transcriptional programs sustain this basal, undifferentiated state, gene-regulatory network inference (SCENIC) was applied across keratinocyte clusters (Aibar et al., 2017). In ASK, the TP63 and PITX1 regulons showed strong selective activity relative to other clusters, and both transcription factors were concordantly upregulated in ASK (**Figure 3E, 1D**), prompting focused analysis of this axis. Because p63 isoforms exert distinct functions – with ΔNp63 maintaining epithelial stemness and opposing Notch-driven differentiation, whereas TAp63 is more linked to differentiation/stress programs (Keyes et al., 2005; Moses et al., 2019; Nguyen et al., 2006) – isoform usage was resolved from 5’ data. *TP63* expression was highest in ASK (p-adj<10⁻J², log_₂_FC=1.65) (**Figure 3F**), and isoform-aware counting demonstrated preferential ΔNp63 expression in ASK compared with basal1/basal2 (p-adj<10⁻¹J), yielding a right-shifted ΔN:TA ratio (**Figure 3G-H**). Consistent with ΔNp63-mediated dampening of differentiation cues, *HES1*, a Notch target involved in keratinocyte differentiation (Moriyama et al., 2006, 2008), showed reduced transcription factor activity (**Figure 3E**), was strongly downregulated in ASK (p-adj < 10⁻JJ, log_₂_FC = -2.26) (**Figure 3J**), and Notch Hallmark signaling scored lowest among keratinocyte clusters (p-adj < 10⁻¹J) (**Figure 3K**). PITX1, a reported cSCC stem-cell factor that cooperates with p63/SOX2 and antagonizes KLF4-driven differentiation (J. Li et al., 2012; Sastre-Perona et al., 2019; Siegle et al., 2014), was likewise SCENIC-active and overexpressed in ASK (**Figure 3E and 1D**). In line with these findings, the AK-derived PM1 keratinocyte line showed higher p63 and PITX1 protein expression than normal human keratinocytes (**Supplementary Figure 3C**) (Hassan et al., 2019).

Finally, consistent with a p63-linked metabolic program in squamous carcinomas (Hsieh et al., 2019), ASK displayed strong upregulation of *SLC2A1*/GLUT1 (p-adj < 10⁻JJ, log_₂_FC = 2.14) with glycolysis pathway enrichment (**Figure 3L; Supplementary Figure 3D-E**). External datasets supported a broader role for p63 in squamous cancers: *TP63* was frequently amplified and positively correlated with *SLC2A1* (GLUT1) across TCGA squamous carcinomas (**Supplementary Figure 3F-G**), and TP63 dependency was enriched in squamous lineages in DepMap (Arafeh et al., 2025); moreover, TP63 knockout caused significant loss-of-fitness in an SCC genome-scale CRISPR screen (FDR 0.00375) (A. L. Ji et al., 2020; Oughtred et al., 2021). Together, the data implicate a ΔNp63/PITX1 regulatory module that maintains ASK in a basal-like, undifferentiated state by attenuating Notch/HES1 signaling and biasing toward glycolytic metabolism.

### 4. ASK gene signature links early AK to invasive cSCC and highlights IGFBP6

An ASK gene signature was derived by differential expression in ASK versus all other keratinocyte subclusters (FDR<0.01; log_₂_FC≥0.5) (**Figure 4A**). A complementary downregulated set (FDR<0.01; log_₂_FC≤-0.5) was generated for sensitivity analyses (**Supplementary Figure 4A**). In the independent spatial transcriptomics cohort with pathologist-annotated regions, the ASK signature was significantly enriched in dysplastic epidermis and depleted from morphologically normal epidermis (**Figure 4B**). Sensitivity analysis using the downregulated gene set showed reciprocal behavior (**Supplementary Figure 4B**). These spatial data confirm the ASK signature in the premalignant keratinocyte compartment.

**Figure 4.**
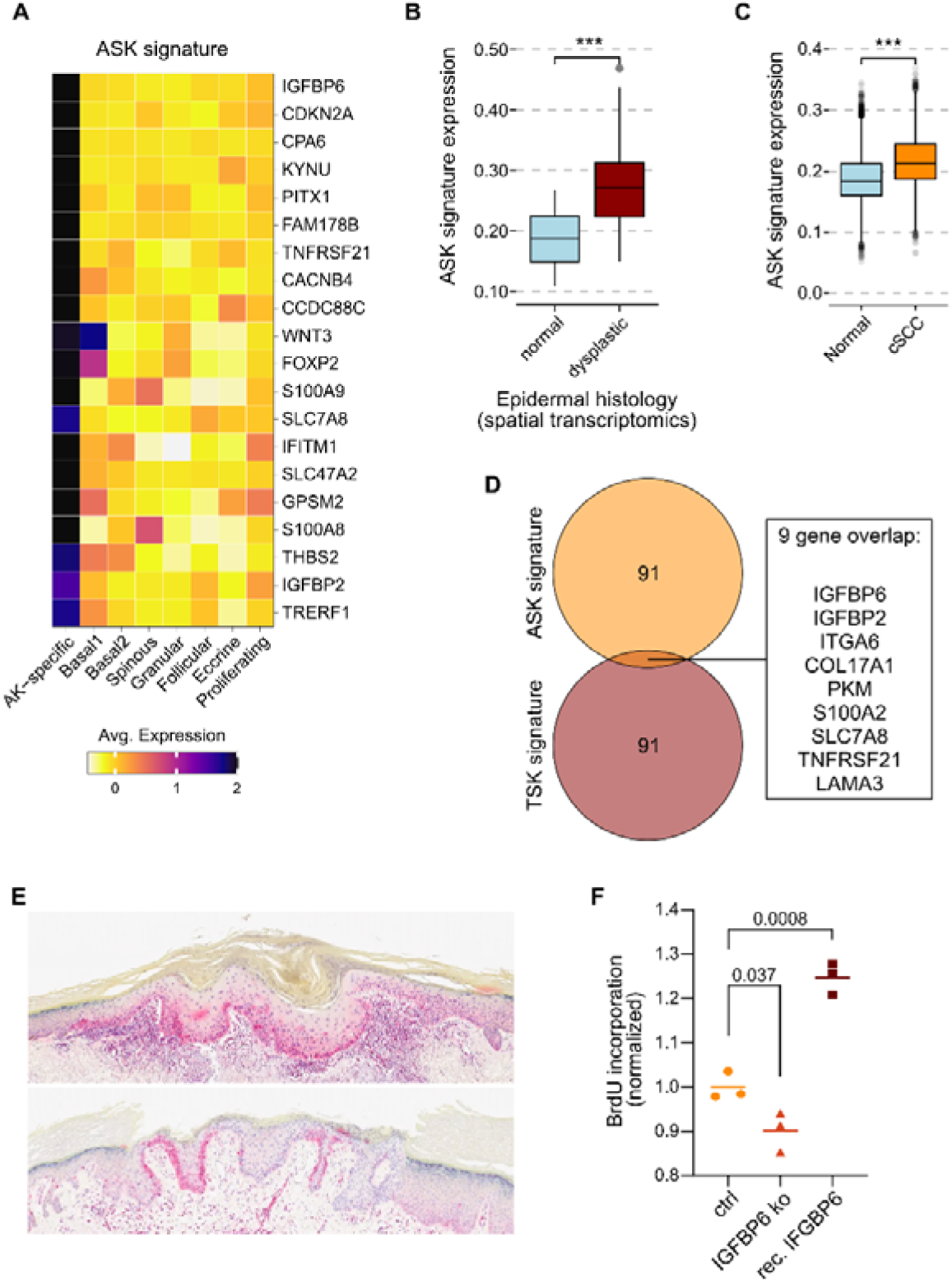
**(A)** Heatmap with average expression of the top upregulated ASK signature genes across keratinocyte subclusters. **(B)** ASK signature enrichment scores in spatial transcriptomics spots annotated as morphologically normal or dysplastic epidermis (Wilcoxon rank-sum test). **(C)** ASK signature enrichment scores in normal and cSCC keratinocytes from Ji et al. (A. L. Ji et al., 2020) (Wilcoxon rank-sum test). **(D)** Venn diagram showing overlap between the ASK and TSK signatures (A. L. Ji et al., 2020). **(E)** IGFBP6 immunohistochemistry on AK tissue. **(F)** BrdU incorporation in PM1 cells following IGFBP6 knockout or treatment with recombinant IGFBP6, normalized to control (t-test). ***: p<0.001. ASK: AK-specific keratinocytes. TSK: tumor-specific keratinocytes. cSCC: cutaneous squamous cell carcinoma.

Projection of the ASK signature onto published scRNA-seq datasets (Hao et al., 2021; A. L. Ji et al., 2020; D.-D. Zou et al., 2023) identified keratinocytes matching ASK (prediction ≥95th percentile) with increasing prevalence from normal to AK to invasive cSCC, as shown above (**Supplementary Figure 1D-E**). Concordant with a progression along the squamous cancer trajectory, the ASK signature was significantly overexpressed in cSCC tumor versus normal keratinocytes from Ji et al. (**Figure 4C**), whereas the ASK-downregulated set was reduced (**Supplementary Figure 4C**) (A. L. Ji et al., 2020).

To relate ASK to invasive cSCC programs, overlap analysis between the ASK and tumor-specific keratinocyte (TSK) gene signatures published by Ji et al. (A. L. Ji et al., 2020) revealed nine shared genes: *IGFBP6, IGFBP2, ITGA6, COL17A1, PKM, S100A2, SLC7A8, TNFRSF21, LAMA3*; while canonical invasion-associated TSK markers (*MMP1, MMP10, PTHLH*) were not elevated in ASK (A. L. Ji et al., 2020) (**Figure 4D**). *IGFBP2* expression had previously been reported in AK (D.-D. Zou et al., 2023). Importantly, the specific TSK markers *MMP1, MMP10* (both implicated in extracellular matrix disassembly) and *PTHLH* (A. L. Ji et al., 2020), were not overexpressed in ASK. *IGFBP6* was among the most upregulated markers of ASK, persisted along the AK to invasive cSCC trajectory, was overexpressed in the AK-derived PM1 keratinocyte cell line (**Supplementary Figure 4D**) (Hassan et al., 2019; Inman et al., 2018) and finally found to be expressed at the protein level by immunostaining of AK (**Figure 4E**). Knockout of *IGFBP6* in PM1 reduced cell proliferation (BrdU incorporation), whereas recombinant IGFBP6 increased proliferation (**Figure 4F**). Together, these data position the ASK signature as an early squamous cancer program detectable in premalignant epithelium that connects to, but is distinct from the invasive TSK program, and nominate *IGFBP6* as a pro-proliferative effector within the ASK compartment.

### 5. Shifts in differentiation and inflammation remodel the AK micro-environment

AK lesions showed a redistribution of the keratinocyte compartment with expansion of an inflammatory basal2 state (33.3% in AK vs. 17.85% in UVES) and near-loss of terminally differentiated granular cells (**Figure 1C**). Basal2 cells expressed inflammatory genes like *IL20*, *CCL2*, and *CXCL2*, and showed high similarity to the AK-cluster described by Li et al. (J. Li et al., 2023) (**Supplementary Figure 5A-B**). Gene-set enrichment supported a global shift toward cytokine/inflammatory programs with concurrent downregulation of epidermal differentiation/barrier pathways (e.g., keratin 1 and 2, filaggrin, transglutaminase 1) (**Figure 5A-B**). Cell–cell communication analysis (CellChat) identified keratinocytes as the principal hub with the largest increase in both outgoing and incoming interactions in AK compared to UVES, exceeding changes in other lineages (**Figure 5C**). Together, these data indicate heightened epithelial inflammation alongside impaired keratinocyte differentiation within AK lesions.

**Figure 5.**
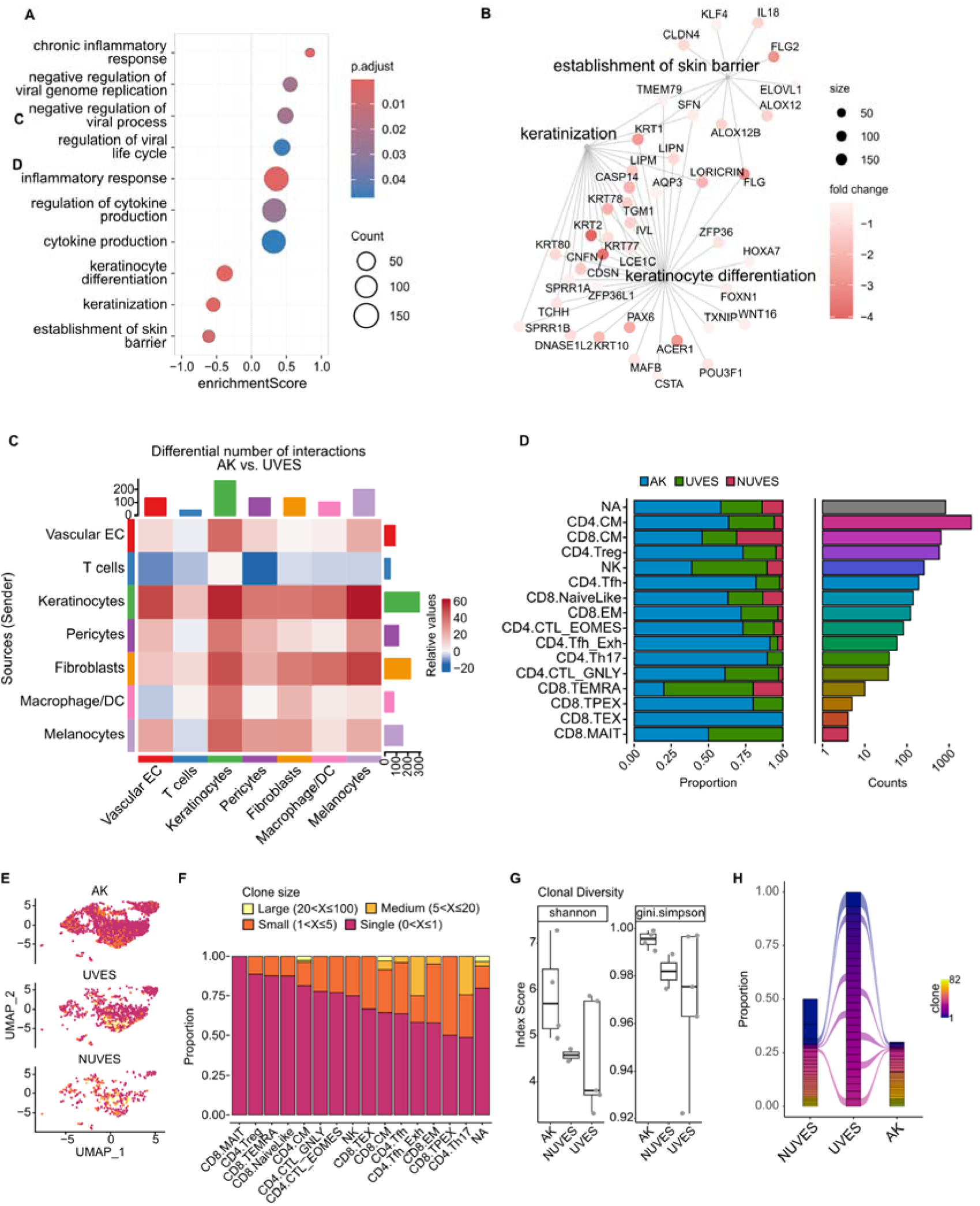
**(A)** Gene set enrichment analysis of differentially expressed genes in AK versus normal skin. Enrichment score sign indicates direction (positive: upregulated in AK; negative: downregulated). **(B)** Gene network plot of downregulated gene sets in AK, grouped by biological process. **(C)** Differential cell-cell interaction heatmap (CellChat) comparing AK to UVES. Values reflect the change in number of inferred interactions per sender-receiver pair. **(D)** Proportion and absolute counts of T cell subclusters by condition. **(E)** UMAP of T cells colored by clone size, shown by condition. **(F)** Clone size distribution as a proportion of each T cell subtype. **(G)** Shannon and Gini-Simpson clonal diversity indices by condition. **(H)** Shared clone distribution across patient-matched AK, UVES, and NUVES. Color indicates clone identity; line thickness reflects clone size. EC: endothelial cell. DC: dendritic cell.

Immune composition mirrored these shifts: lymphocytes increased in AK (t-statistic > 1.8, p<0.1), with the most abundant subtypes being CD4+ and CD8+ central memory (CM) and CD4+ regulatory T cells (CD4.Treg) (**Figure 5D**). In AK, T cell exhaustion and suppression was detected: CD8+ exhausted T-cells (CD8.TEX) were found only in AK, and proportions of CD4+ T follicular helper cells (CD4.Tfh_Exh) and CD4.Tregs increased (t-statistic > 1.21, AK vs. UVES) (**Figure 5D**).

Paired single-cell T-cell receptor (TCR) sequencing showed predominantly polyclonal repertoires with some oligoclonal expansions (**Figure 5E-F**), especially within CD4.Th17 and CD4.Tfh_Exh. Analysis of the TCR repertoire showed a non-significant trend for higher clonal diversity in AK compared to normal skin (**Figure 5G**), and some shared clones across patient matched sites, consistent with recall responses to common viral antigens (**Figure 5H; Supplementary Figure 5C-E**).

Taken together, the disrupted keratinocyte differentiation cascade and chronic inflammatory activation create a barrier-deficient, inflamed niche that may support persistence and progression of epidermal dysplasia.

## Discussion

This study provides a multimodal, patient-matched characterization of AK at single-cell resolution, integrating simultaneous transcriptomic and surface-proteomic profiling with spatial whole-transcriptome analysis. The central finding is the identification of AK-specific keratinocytes (ASK): a genomically malignant, transcriptionally undifferentiated keratinocyte population within the dysplastic epithelial layer, governed by a ΔNp63/PITX1 regulatory axis, and transcriptionally related to – yet distinct from - the tumor-specific keratinocytes (TSK) of invasive cSCC.

### An early, genomically defined malignant keratinocyte population

The ASK population expands on recent single-cell AK studies that reported disease-associated keratinocyte states and upregulation of *CDKN2A* and *IGFBP2* (J. Li et al., 2023; D.-D. Zou et al., 2023) by providing further genomic support for pre-malignant identity. InferCNV revealed recurrent copy number alterations characteristic of early squamous carcinogenesis, specifically loss of chromosome 9p (*CDKN2A*) and gain of 8q (*MYC*), in ASK (Kim, Jung, et al., 2023; Thomson et al., 2021). Somatic variant calling identified a dominant UV-associated SBS7b signature and the highest mutational burden across all keratinocyte clusters in ASK, placing it between UV-exposed normal skin and invasive cSCC in the expected mutational trajectory (Pickering et al., 2014; Thomson et al., 2021). The paradoxical co-occurrence of *TP53* overexpression with reduced p53 pathway activity and *CDKN1A* downregulation is consistent with dominant-negative *TP53* mutations that result in p53 protein accumulation, commonly identified in AK (Blokx et al., 2003; Nelson et al., 1994; Sung et al., 2022). Such mutations accumulate in UV-exposed skin well before histological dysplasia is apparent and are considered among the earliest drivers of clonal keratinocyte expansion. Spatial deconvolution confirmed ASK to the dysplastic basal layer, absent from morphologically normal epidermis, supporting a model in which UV-driven p53 inactivation permits outgrowth of a genomically unstable progenitor population that avoids terminal differentiation from which further progression can arise (Klein et al., 2010; Ziegler et al., 1994).

### A **Δ**Np63/PITX1 regulatory module maintains an undifferentiated state

Gene regulatory network inference identified selective *TP63* and *PITX1* regulon activity as key transcriptional determinants of ASK identity. Isoform-resolved analysis confirmed predominance of the ΔNp63 isoform, consistent with its established role as a master regulator of epithelial progenitor maintenance and an oncogenic driver in squamous carcinomas (Gatti et al., 2019; Moses et al., 2019). p63 selectively suppresses *HES1* transcription to attenuate Notch-driven differentiation and maintain an undifferentiated basal state in keratinocytes (Nguyen et al., 2006), consistent with the strong *HES1* downregulation and markedly reduced Notch hallmark activity observed in ASK. This relationship is bidirectional: Notch1 also suppresses ΔNp63 in normal epidermis to promote differentiation commitment (Nguyen et al., 2006), meaning that the frequent early *NOTCH1* loss-of-function mutations in squamous carcinogenesis (South et al., 2014) would de-repress ΔNp63, further amplifying the undifferentiated phenotype observed here. PITX1 is a *de novo* expressed transcription factor in cSCC tumor-propagating cells (Sastre-Perona et al., 2019; Siegle et al., 2014) that promotes self-renewal while antagonizing KLF4-driven differentiation through a feed-forward circuit with p63 and SOX2 (Sastre-Perona et al., 2019). Its co-activity with ΔNp63 in ASK, together with the marked reduction of KLF4 regulon activity, is consistent with PITX1-mediated suppression of keratinocyte differentiation. The present data thus place this co-regulatory module as early as the AK stage, suggesting that the transcriptional circuitry associated with squamous cancer stem cell identity is already operative in pre-invasive lesions. Downstream of this axis, ASK displayed marked *SLC2A1*/GLUT1 upregulation and activation of glycolytic gene programs. The positive correlation between *TP63* and *SLC2A1* expression across TCGA squamous carcinoma datasets, together with *TP63* dependency data from DepMap and a cSCC CRISPR screen, further support p63-driven glycolytic reprogramming as a lineage-selective dependency established prior to invasion, consistent with direct transcriptional activation of *SLC2A1* by p63 and SOX2 demonstrated in squamous carcinoma models (Hsieh et al., 2019).

### Shared dependencies and distinct invasion programs in ASK and invasive cSCC

Overlap analysis between the ASK and TSK signatures (A. L. Ji et al., 2020) revealed nine shared genes (*IGFBP6, IGFBP2, ITGA6, PKM, LAMA3*, and others) representing core oncogenic dependencies rather than invasion markers. In contrast, canonical TSK effectors of matrix proteolysis and paracrine invasion signaling, namely *MMP1*, *MMP10*, and *PTHLH*, were absent from ASK (Pittayapruek et al., 2016; Takayama et al., 2010). This pattern supports a two-phase model: an early phase (ASK) in which core dependencies are established while the invasive program is absent, and a later phase (TSK) in which MMP and PTHLH expression licenses stromal penetration. The progressive increase of ASK-like cells from normal skin to AK to invasive cSCC in independent published datasets further suggests that this population persists and expands throughout carcinogenesis rather than representing a transient intermediate, implying that molecular targets operative in ASK may remain relevant in established tumors.

Among shared markers, *IGFBP6* was among the highest-ranked ASK genes, confirmed at the protein level in AK tissue, and tracked along the normal-to-cSCC trajectory. Functional experiments in the PM1 AK-derived cell line showed reduced proliferation upon *IGFBP6* knockout and increased proliferation with recombinant protein. These in vitro data are preliminary, and the mechanisms by which IGFBP6 influences keratinocyte proliferation in this context warrant further investigation, but nominate IGFBP6 as a potential pro-proliferative effector.

### Surface markers and microenvironmental context

CITE-seq identified elevated CD49f (*ITGA6*) and CD26 (*DPP4*) on ASK at both protein and RNA levels. CD49f marks keratinocyte stem cells and tumor-initiating populations in cSCC (Schober & Fuchs, 2011), and its expression in ASK is consistent with their basal-like progenitor identity. Elevated CD26/DPP4 activity has been reported in AK lesional skin (Moehrle et al., 1995), and DPP4 inhibition reduces keratinocyte tumor growth in murine models (Arwert et al., 2012) and promotes epidermal differentiation in organotypic cultures (Bao et al., 2022). Given the undifferentiated phenotype and high DPP4 surface expression of ASK, DPP4 activity may contribute to maintaining the differentiation block in this population, and its pharmacological inhibition could help restore normal differentiation trajectories in addition to reducing proliferation. Consistent with this, DPP4 inhibitor use in type 2 diabetes patients has been associated with a reduced incidence of skin neoplasms (Z. Li et al., 2022), raising the possibility of chemopreventive activity in high-risk patients with extensive AK field cancerization.

Beyond ASK, AK lesions showed expansion of an inflammatory basal keratinocyte state expressing *IL20*, *CCL2*, and *CXCL2*, and depletion of granular cells, indicating a global shift toward barrier disruption and chronic epidermal inflammation consistent with the disease-associated cluster described by (J. Li et al., 2023; D.-D. Zou et al., 2023). CellChat analysis identified keratinocytes as the principal hub of increased cell-cell communication in AK, suggesting the epithelial compartment actively shapes the permissive niche rather than being a passive recipient of mutagenic damage. In the immune compartment, T cell infiltration increased in AK with a shift toward memory phenotypes, the appearance of exhausted CD8+ T cells exclusively in lesional tissue, and expansion of CD4+ Tregs and Th17-skewed populations with oligoclonal TCR expansions. The Th17 skewing is consistent with UV-driven expansion of IL-17-producing CD4+ T cells in murine skin tumors, where IL-17 has been shown to promote tumor growth (Nasti et al., 2011). Together, these findings indicate that immunosuppressive and tumor-promoting immune circuits are already being established at the pre-invasive stage, a pattern that intensifies with cSCC grade (D.-D. Zou et al., 2023).

### Conclusions

In conclusion, using CITE-seq and spatial transcriptomics we identified a discrete population of malignant keratinocytes in AK, marked by UV mutational burden, copy number alterations, a basal-like undifferentiated phenotype, and ΔNp63/PITX1 regulatory activity, that shares core gene networks with tumor-propagating cells in invasive cSCC while lacking invasion-associated effectors. These findings define the molecular identity of early malignant keratinocytes and position IGFBP6, CD26, and CD49f as candidate markers of the pre-invasive malignant compartment.

Since AK represents the earliest histologically defined stage of squamous carcinogenesis, molecular programs already active at this pre-invasive stage and sustained through progression are likely to represent core dependencies of established tumors. Targeting such programs may prove more difficult for tumor cells to escape under therapeutic pressure than targeting alterations acquired later, and could thus inform treatment strategies for advanced cSCC. Finally, AK offers a distinct practical advantage as a model of squamous carcinogenesis: it is minimally invasively accessible and amenable to longitudinal study in vivo. Given the shared lineage dependency on p63 across squamous carcinomas of the skin, lung, head and neck, and esophagus (Campbell et al., 2018; Dotto & Rustgi, 2016), insights derived here may be relevant beyond cSCC to a broader group of epithelial malignancies that collectively represent a major cause of cancer morbidity and mortality worldwide.

## Methods

### Primary human tissue and single-cell library preparation

Skin biopsies were collected from consenting patients and each sample was halved. One half was formalin-fixed and paraffin-embedded (FFPE) and used for histological confirmation by a board-certified dermatopathologist (I.K.). The second half was immediately live slow-frozen by the Dermatology Biobank of the University Hospital Zurich, to be used for single-cell RNA sequencing, as previously described (Restivo et al., 2022). The tissue was cut in 2×2 mm pieces and submerged in 1 ml freezing medium (90% FBS (Biowest, cat. no. S006420E01, batch no. S169419181H) with 10% DMSO (Sigma, cat. no. 102148154)) in a cryovial (Sarstedt, cat. no. 72.380.002), followed by slow-freezing in a CoolCell® Container (Corning, cat. no. 432001) in a -80 °C freezer.

#### CITE-seq

Live slow-frozen biobanked tissue samples were processed for CITE-seq as previously described (Burja et al., 2022; Restivo et al., 2022). Samples were quickly thawed in a water bath, re-suspended in 10 ml cold PBS with 0.04% BSA (Sigma-Aldrich, cat. no. A7906) and incubated for 10 min to allow DMSO diffusion from the cells. Samples were centrifuged (300 g, 5 min) and cut into small pieces using scalpels. Enzymatic dissociation was performed in three steps, according to the protocol by Haertel et al. (Haertel et al., 2018), which was optimized for single-cell analysis of surface proteins from skin tissue. First, small tissue pieces were incubated in 5 ml of Mg^2+^/Ca^2+^-free PBS with 10 mM EGTA for 20 min at 37 °C on an orbital shaker at 100 rpm, then centrifuged (300 g, 5 min) and washed with PBS. Second, the tissue was incubated in 2 ml RPMI 1640/HEPES with 1.3 U/m Liberase TL (Roche, cat. no. 5401020001) and 15 KU DNase I (Sigma, cat. no. D5025) at 37°C for 60-90 min with magnetic stirrers at slow speed. Third, 1 KU/ml Dispase II (Roche, cat. no. 04942078001) and 7.5 Mm MgCl_2_ were added. Samples were incubated for 15 min at 37 °C, followed by 15 min at room temperature. Cells were centrifuged (300 g, 5 min), washed with PBS with 0.04% BSA, and passed through a 100 µm cell strainer (Falcon, cat. no. 352360) and a 35 µm cell strainer (Falcon, blue capped FACS tubes, cat. no. 352235). Cell count and viability was accessed on Luna-FL^TM^ cell counter (Dual Fluorescence Cell counter, Logos Biosystems Inc., cat. no. L1001) using Acridine Orange/Propidium Iodide live/dead staining (AOPI, Logos Biosystems Inc., cat. no. F23001) with counting slides (PhotonSlides, ultra-low fluorescence counting slides, Logos Biosciences Inc., cat. no. L12005). Cells were resuspended in PBS with 1% BSA for staining following the demonstrated protocol for Cell Surface Protein Labeling from 10x Genomic (*10xgenomics.com*, n.d.). Briefly at least 10^5^ cells were incubated with Human TruStain FcX (BioLegend, cat. no. 422301). After Fc receptor blocking, the TotalSeq™-C Human Universal Cocktail, V1.0 (BioLegend, cat. no. 399905) antibody panel was added and samples were incubated for 30 min on ice. Finally, cells were washed four times, resuspended in PBS with 1% BSA, assessed for viability and adjusted for optimal cell concentration according to 10x Genomics recommendations (700-1200 cells/µl).

Single-cell droplets were generated using the Chromium Next GEM Single Cell 5’ kit v2 (Dual Index) (10x Genomics, cat. no. 1000263), a 10x Genomics Chromium Single Cell Controller (10x Genomics, cat. no. PN110211), and Chromium Next GEM Chip K (10x Genomics, cat. no. PN2000182), according to the manufacturer instructions. Paired-end sequencing (PE 26/10/10/90) was performed on an Illumina NovaSeq SP flow cell. According to the recommendations of 10x Genomics, a sequencing depth of >20,000 read pairs per cell for gene expression libraries and >5,000 read pairs per cell for surface protein libraries was ensured.

#### scTCR-seq

TCR libraries were prepared from amplified cDNA by nested PCR using the Chromium Single Cell Human TCR Amplification Kit (10x Genomics, cat. no. 1000252). TCR pools were adjusted based on expected cell numbers and PE 151/8/0/151 sequenced using Illumina NovaSeq with a targeted coverage of 5,000 read pairs per cell, according to 10x Genomics recommendations for V(D)J.

### CITE-seq analysis

Raw transcriptomic reads were demultiplexed and aligned against the human reference genome assembly GRCh38.p13 using the 10x Genomics CellRanger v6.0.2 pipeline (Zheng et al., 2017). The R package Seurat v5.0.2 (Hao et al., 2021) was used for the downstream analyses of the filtered count matrices. Cells with unique feature counts <250 or >4,000-6,000, unique UMI counts >20,000, mitochondrial gene counts >15-30% and ribosomal gene counts >40% were discarded as part of quality control. Filtered samples were log normalized and integrated using canonical correlation analysis (CCA). Integrated data was scaled and principal component analysis (PCA) was performed using the top 2,000 variable features for dimensional reduction. Samples were clustered together using the Louvain algorithm. For each cell cluster, Wilcoxon rank-sum test was applied to identify the marker genes. Adjusted p-values < 0.05 were considered significant. Cell clusters were annotated based on known markers from literature (Restivo et al., 2022; Solé-Boldo et al., 2020; Z. Zou et al., 2021). For testing differences in cell type proportions, the function *propeller()* from the package speckle v0.99.7 was used (Phipson et al., 2022).

**Signature enrichment scores** were calculated using the UCell package v2.4.0 (Andreatta & Carmona, 2021). Statistical significance was calculated using the Wilcoxon rank-sum test.

### Weighted-nearest neighbor integration of surface protein expression

Surface protein expression was normalized against the background ambient noise and cell-to-cell technical noise by combining the isotype control levels using the dsb package v1.0.2 (Mulè et al., 2022). Normalized protein expression was added as a separate assay to the filtered cells. PCA was separately performed using only the surface proteins for dimensional reduction. For each cell, closest neighbors were calculated based on the top assay specific principal components using a weighted combination of RNA and surface protein similarities. Cells were integrated and clustered together based on their multimodal neighbors and cell specific modality weights using the Seurat v5.0.2 (Hao et al., 2021) pipeline.

#### Keratinocyte subclustering

The keratinocyte cluster was subset and doublets containing T cells (PTPRC, CD3E) (<1% of keratinocytes), endothelial cells (PECAM1) (<4%) and smooth muscle cells (ACTA2) (<0.7%) were removed, followed by SCTransform v2 normalization (Choudhary & Satija, 2022; Hafemeister & Satija, 2019) and reclustering. Differential gene expression analysis was performed with FindMarkers() function from the Seurat package v5.0.2.

**Gene set enrichment analysis** was performed and visualized with the clusterProfiler v4.8.3 (Wu et al., 2021) and the enrichplot v1.20.3 (*Enrichplot*, n.d.) packages using Biological Processes (BP) terms from Gene Ontology.

**Copy number aberrations** were inferred with the infercnv package v1.16.0 (*Infercnv: Inferring CNV from Single-Cell RNA-Seq*, n.d.). The keratinocytes in NUVES were used as reference cells. For visualization of results, the SCPubr package v2.0.2 package and the do_CopyNumberVariantPlot() function (Blanco-Carmona, 2022) was used.

**Somatic single-nucleotide variants** were called directly from scRNA-seq data using SComatic v1.0 (Muyas et al., 2024). BAM files generated by CellRanger were used as input together with cell-type annotations from the Seurat clustering. Variants were called per cell-type cluster using the default SComatic pipeline. Only variants supported by ≥3 cells within a cluster and absent from NUVES keratinocytes used as the germline reference were retained. Mutational load was calculated as the number of somatic single-nucleotide variants per megabase of transcribed sequence per keratinocyte subcluster. The resulting variant call sets were formatted as VCF files and uploaded to the web-based SigProfilerAssignment tool (Díaz-Gay et al., 2023) for decomposition into COSMIC single-base substitution (SBS) signatures v3.3 against the GRCh38 reference. Reconstruction accuracy was assessed by cosine similarity between observed and reconstructed mutational profiles.

#### TP63 isoform calling

TP63 isoform-specific read counts were derived from 5’ CITE-seq BAM files. Gene annotation was imported from the reference GTF file and filtered to TP63 exons. Isoform-distinguishing exons were defined as: for TAp63, exons ENSE00001343828.2, ENSE00003502961.1, and ENSE00001184832.1 (absent from ΔNp63); for ΔNp63, exon ENSE00001710939.1 (absent from TAp63). Per-cell deduplicated reads overlapping each exon were counted using the ezReadGappedAlignments() function from the ezRun package and added to the Seurat object as cell-level metadata to compute per-cell ΔN:TA ratios and visualize isoform distributions across keratinocyte subclusters.

**Gene Regulatory Network (GRN) analysis** was performed for keratinocyte subclusters using the R package SCENIC v1.2.4 (Aibar et al., 2017). Potential transcription factor targets were detected based on the filtered gene expression matrix. Once the GRN was built on co-expression modules, network activity was analyzed in the keratinocytes via regulon scores.

**Pairwise mean squared error (MSE) distances**, a robust measure of similarity in single cell data (Y. Ji et al., 2023), between keratinocyte subclusters were calculated using a custom R function mse_distance and the future_lapply function for parallel computation. The resulting distances were visualized as a heatmap using the ComplexHeatmap package v2.16.0 (Gu et al., 2016). A Monte Carlo analysis with 100 iterations using 80% of the data was also conducted to assess the robustness of the findings. The median distance for each subcluster was computed and plotted as a violin plot using the ggplot2 package.

#### T cell annotation

T and NK cells were subset and UMAP was recomputed. The scGate package (Andreatta, Berenstein, et al., 2022) was used to classify cells as CD4+ or CD8+ T cells. The ProjecTILs package was then used to classify the T cells into functional clusters using the *ProjecTILs.classifier()* function (Andreatta et al., 2021). The CD8 and CD4 T cell reference maps were downloaded from the Swiss Portal for Immune Cell Analysis (SPICA) (Andreatta, David, et al., 2022). The results were plotted using the DimPlot and VlnPlot functions from the Seurat package.

#### TCR repertoire analysis

Raw V(D)J reads were processed using CellRanger vdj (Zheng et al., 2017). Clone size, shared clones, and clonal diversity were analyzed using the scRepertoire package v2.0.0 (Borcherding et al., 2020). Cell-level clonotype data were mapped to the Seurat objects containing cell typing and gene expression information. Clonotypes were determined by V(D)J-recombinations, i.e. in case of identical TCR α-and β-chains and the same CDR3 amino acid, cells were defined as clonal using the *combineExpression()* function. Matching of TCR sequences to known antigens was done using the Trex package (Mudd et al., 2023).

#### CellChat

Intercellular communication was inferred, analyzed and visualized using the CellChat package v1.6.1 (Jin et al., 2021). Cell types with >100 cells in each condition were included. Secreted signaling from CellChatDB was used as the ligand-receptor interaction database. Only cell-cell interactions occurring in at least 10 cells were included in the analysis. The netVisual_heatmap() was used for visualizing the cellular interactions.

### Spatial transcriptomics

From the Dermatology Biobank at the University Hospital Zurich, FFPE blocks with AK were selected by a board-certified dermatopathologist (I.K.) based on representative histology and available tissue material in the block. From two consecutive 10 µm cuts, RNA was extracted using Deparaffinization Solution (QIAGEN, cat. no. 19093) and the RNeasy FFPE Kit (QIAGEN, cat. no. 73504). RNA fragmentation was assessed by determination of DV200 using the Agilent TapeStation 4200 with the RNA ScreenTape (Agilent, cat. no. 5067-5576) and RNA buffer (Agilent, cat. no. 5067-5577). As per 10x Genomics recommendations, only samples with DV200 >30% were processed for Visium CytAssist.

Raw reads were demultiplexed and aligned using 10x SpaceRanger, followed by tissue and fiducial detection and barcode/UMI counting. Through the Loupe browser v7.0 (10x Genomics), regions of dysplastic and morphologically normal epidermis were annotated by a board-certified dermatopathologist (I.K.). With the Seurat package v5.0.2 (Hao et al., 2021), the spot by gene expression data was normalized using sctransform v2 (Hafemeister & Satija, 2019). Cell type deconvolution was performed via integration of the scRNA-seq dataset following the following the Seurat spatial vignette (https://satijalab.org/seurat/articles/spatial_vignette).

### Immunohistochemistry

IGFBP6 immunohistochemistry on FFPE sections of AK tissue was performed on a Ventana BenchMark Ultra automated platform (Roche) with the UltraView Universal Alkaline Phosphatase Red Detection Kit (Roche, cat. no. 05269814001) using anti-IGFBP6 antibody (Sigma-Aldrich, cat. no. HPA008005) at 1:200 dilution according to the manufacturer’s instructions.

### Cell culture and functional experiments

The pre-malignant AK-derived keratinocyte cell line PM1 (Hassan et al., 2019) was cultured in keratinocyte serum-free medium (K-SFM) supplemented with epidermal growth factor and bovine pituitary extract at 37 °C in a humidified atmosphere with 5% CO_₂_. Normal human primary keratinocytes (HPK) were isolated from human foreskin as previously described (Grossi et al., 2020).

#### Western blot

Cell lysates were harvested in SDS loading buffer containing 0.1 M Tris-HCl pH 6.8 and 4% SDS, supplemented with 0.1 M DTT as reducing agent. Lysates were incubated at 95 °C for 5 minutes and sonicated. Proteins were resolved by SDS-PAGE. Following primary antibodies were used: anti-p63 (D9L7L, Cell Signaling Technology, cat. no. 39692, 1:1000), anti-PITX1 (ab70273, Abcam, 1:2500), anti-IGFBP6 (ab219560, Abcam, 1:1000), anti-Keratin-10 (B433, 1:1000), anti-Filaggrin (B474, 1:1000), anti-Loricrin (B477, 1:1000), anti-β-actin (E328, 1:5000). Dried membranes were scanned on a LiDE 210 scanner (Canon Inc.). Densitometry was performed using ImageJ and normalized to β-actin.

#### Keratinocyte differentiation assay

As previously described (Fenini et al., 2018), HPK and PM1 cells were seeded at 2×10J cells per well in 12-well plates and grown in K-SFM supplemented with epidermal growth factor and bovine pituitary extract to confluence. Differentiation was induced by switching to KBM-Gold medium (Lonza, cat. no. 10130685) supplemented with gentamicin and hydrocortisone. Cell lysates were collected at days 0, 2, 4, 7, and 10 for western blot analysis of differentiation markers (keratin 10, filaggrin, loricrin).

#### IGFBP6 CRISPR-Cas9 knockout

Guide RNA design was performed using CHOPCHOP (https://chopchop.cbu.uib.no/) with the following settings: target organism Homo sapiens (hg38/GRCh38), CRISPR/Cas9 for knockout, targeting all exons including UTRs, searching exons and immediate short flanking regions, with isoform consensus determined by union of all isoforms. Candidate guide sequences were verified for exon location using Benchling (https://benchling.com) and screened for off-target sites using Off-Spotter (https://cm.jefferson.edu/Off-Spotter/). The guide RNA targeting exon 1 of *IGFBP6* (5’-GGACGACGAGGCGCCTTTGC-3’) was selected. IGFBP6 was knocked out in PM1 cells by CRISPR-Cas9 ribonucleoprotein (RNP) electroporation following the protocol of Karakaya et al. (Karakaya et al., 2024). Briefly, 36 pmol TrueCut HiFi Cas9 protein (ThermoFisher, cat. no. A50575) and 36 pmol guide RNA (TrueGuide custom gRNA, ThermoFisher) were complexed and used to electroporate 1.5×10J PM1 cells on a Neon Transfection System (MPK5000, Invitrogen) with the following settings: 1700 V, 20 ms, 1 pulse. A non-targeting guide RNA served as control. Knockout efficiency was confirmed by western blot.

#### Recombinant IGFBP6 treatment

For recombinant protein experiments, PM1 cells were treated with recombinant human IGFBP6 (gibco, cat. no. 350-07B-20UG) for 5 days at 375 ng/ml, according to published methods (Aboalola & Han, 2017); vehicle-treated cells served as controls.

#### BrdU proliferation assay

Cell proliferation was assessed by BrdU incorporation, using a colorimetric Cell Proliferation ELISA kit (Roche, cat. no. 11647229001), according to the manufacturer’s instructions. BrdU incorporation was quantified measuring absorbance with Tecan Spark 10M plate reader and normalized to the respective control condition. Statistical significance was assessed by t-test.

### Analysis of published datasets

#### Published scRNA-seq datasets

The scRNA-seq datasets GSE193304 (D.-D. Zou et al., 2023) and GSE144236 (A. L. Ji et al., 2020) were accessed via NCBI GEO and analyzed with the Seurat v5.0.1 package in R. Cell type information was transferred from the dataset generated here to the external dataset using the *FindTransferAnchors()* and *TransferData()* functions. The 95th percentile was used as a cut-off for the prediction scores.

#### PM1 cell line expression data

Gene expression data were obtained from the Gene Expression Omnibus (GEO, accession GSE98767 (Inman et al., 2018)) using the GEOquery R package GEOquery v2.68.0 (Davis & Meltzer, 2007). This dataset contains microarray expression profiles of normal human keratinocytes and pre-malignant keratinocyte cell lines. Log2-transformed expression values were extracted for genes of interest. Expression levels were compared between PM1 (pre-malignant) and normal keratinocyte samples using the Wilcoxon rank-sum test.

#### TCGA data

Copy-number and RNA expression data for squamous cell carcinomas of the lung (TCGA-LUSC), head and neck (TCGA-HNSC), and cervix (TCGA-CESC) were queried from cBioPortal (https://www.cbioportal.org). *TP63* amplification frequencies and *TP63–SLC2A1* correlations (assessed by Spearman and Pearson correlation) were extracted as reported by the portal. Tumor versus paired normal expression comparisons were obtained from TNMplot (https://tnmplot.com/analysis/) (Bartha & Győrffy, 2021).

#### Genetic dependency

*TP63* dependency scores and lineage enrichment were retrieved from the DepMap portal (https://depmap.org/portal/) (Arafeh et al., 2025).

#### CRISPR screen

Results of the SCC genome-scale CRISPR screen by Ji et al. 2020 (A. L. Ji et al., 2020) were accessed via the BioGRID Open Repository of CRISPR Screens (ORCS; https://orcs.thebiogrid.org/) (Oughtred et al., 2021).

## Ethics declaration

The collection and use of clinical material for research purposes was approved by the Cantonal Ethics Committee Zurich (BASEC PB.2018-00194, KEK 2017-00688, KEK2019-02150) and patient informed consent was obtained for all human primary material.

## Acknowledgments

We thank the team of the Dermatology Biobank, University Hospital Zurich for assistance with primary samples. Sequencing was performed at the Functional Genomics Center Zurich (FGCZ) of University of Zurich and ETH Zurich. Figure 1A was created with BioRender.com.

PM1 cells were kindly gifted by Irene Leigh and Gernot Walko.

RS, AT, MDF, HDB, ML, and RD are members of the SKINTEGRITY.CH collaborative research program. RS is supported by the Swiss Cancer Research Foundation and the Swiss National Science Foundation (323630_207029).

## Supplementary Figures

**Supplementary Figure 1.**
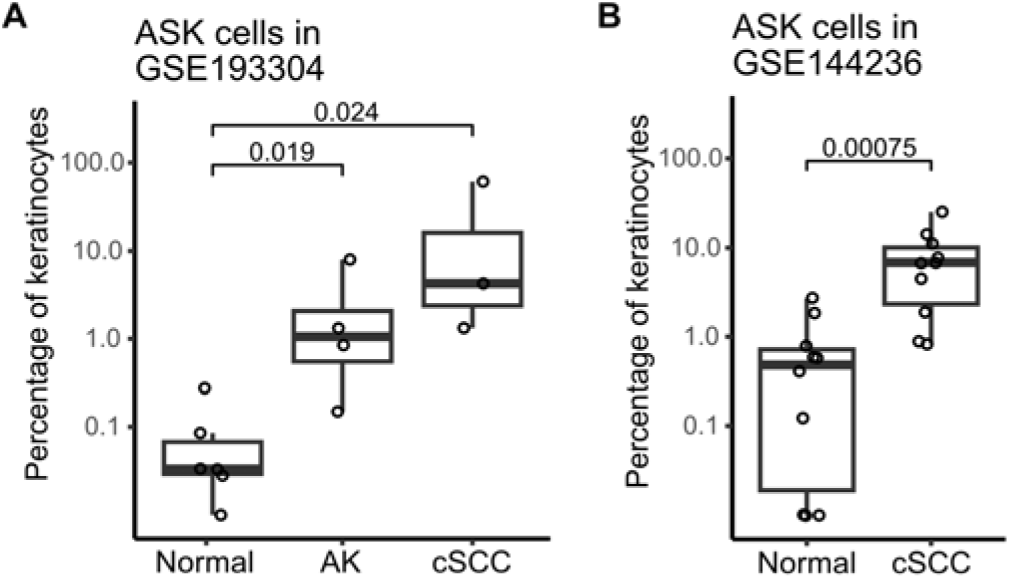
ASK cells are progressively enriched across the squamous carcinogenesis trajectory in independent datasets. **(A)** Percentage of keratinocytes with the highest ASK prediction scores in GSE193304 (D.-D. Zou et al., 2023), by disease stage (Wilcoxon rank-sum test). **(B)** As in (A), for GSE144236 (A. L. Ji et al., 2020). ASK: AK-specific keratinocytes. AK: actinic keratosis. cSCC: cutaneous squamous cell carcinoma.

**Supplementary Figure 2.**
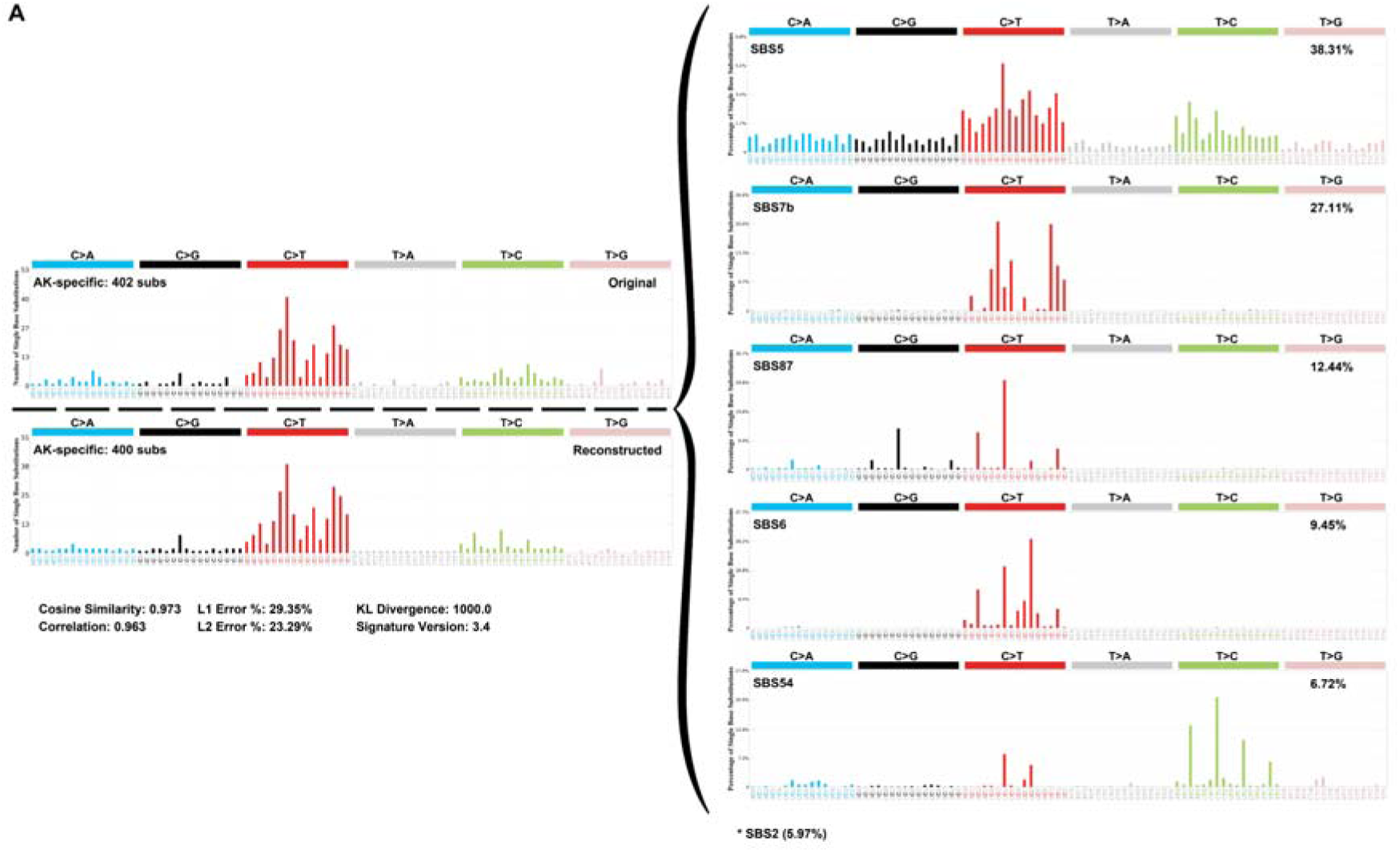
Mutational signature reconstruction for the ASK profile. **(A)** COSMIC single-base substitution (SBS) signature reconstruction for the ASK mutational profile. The observed (original) and reconstructed profiles are shown on the left; contributing signatures and their relative proportions on the right. Overall reconstruction quality: cosine similarity 0.973.

**Supplementary Figure 3.**
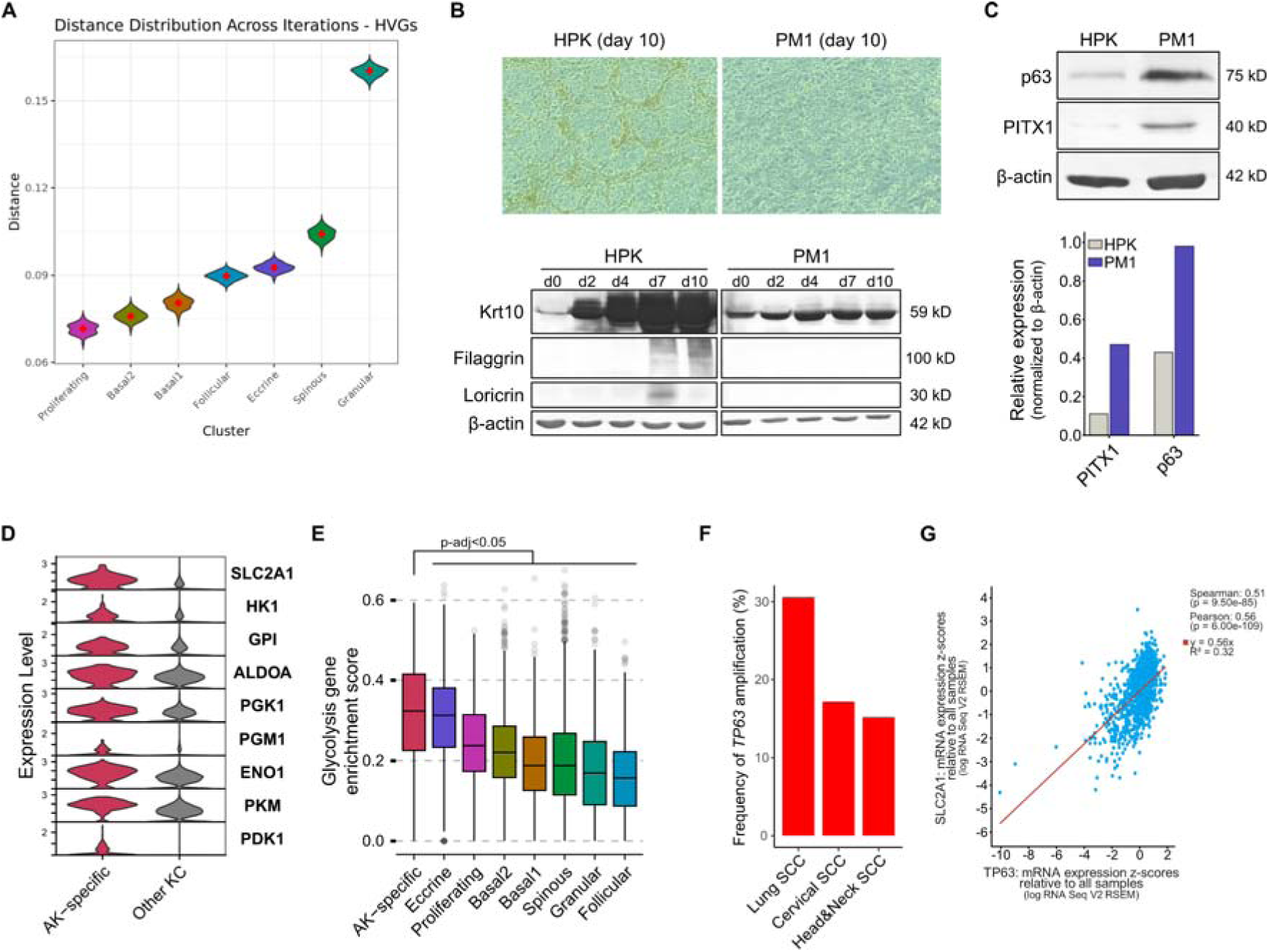
**(A)** Distribution of pairwise mean squared error (MSE) distances between ASK and each remaining keratinocyte subcluster across 100 Monte Carlo iterations (80% subsampling per iteration). Red diamonds indicate median distances. HVG: highly variable genes. MSE: mean squared error. **(B)** Differentiation assay comparing normal human primary keratinocytes (HPK) and the AK-derived PM1 cell line. Top: phase-contrast images at day 10 of culture in differentiation medium. Bottom: western blot analysis of differentiation markers keratin 10 (Krt10), filaggrin, and loricrin at days 0, 2, 4, 7, and 10. β-actin serves as loading control. **(C)** Western blot analysis of p63 and PITX1 protein expression in normal human primary keratinocytes (HPK) and the AK-derived PM1 cell line. β-actin serves as loading control. Bar chart shows densitometry quantification normalized to β-actin. Data from a single experiment. **(D)** Expression of glycolytic genes in ASK versus other keratinocyte subclusters. **(E)** Glycolysis gene set enrichment scores by keratinocyte subcluster (Wilcoxon rank-sum test). **(F)** Frequency of *TP63* amplification across squamous cell carcinomas in TCGA. **(G)** Correlation between *TP63* and *SLC2A1* mRNA expression across lung, cervical, and head and neck SCC in TCGA (Spearman r=0.51, Pearson r=0.56).

**Supplementary Figure 4.**
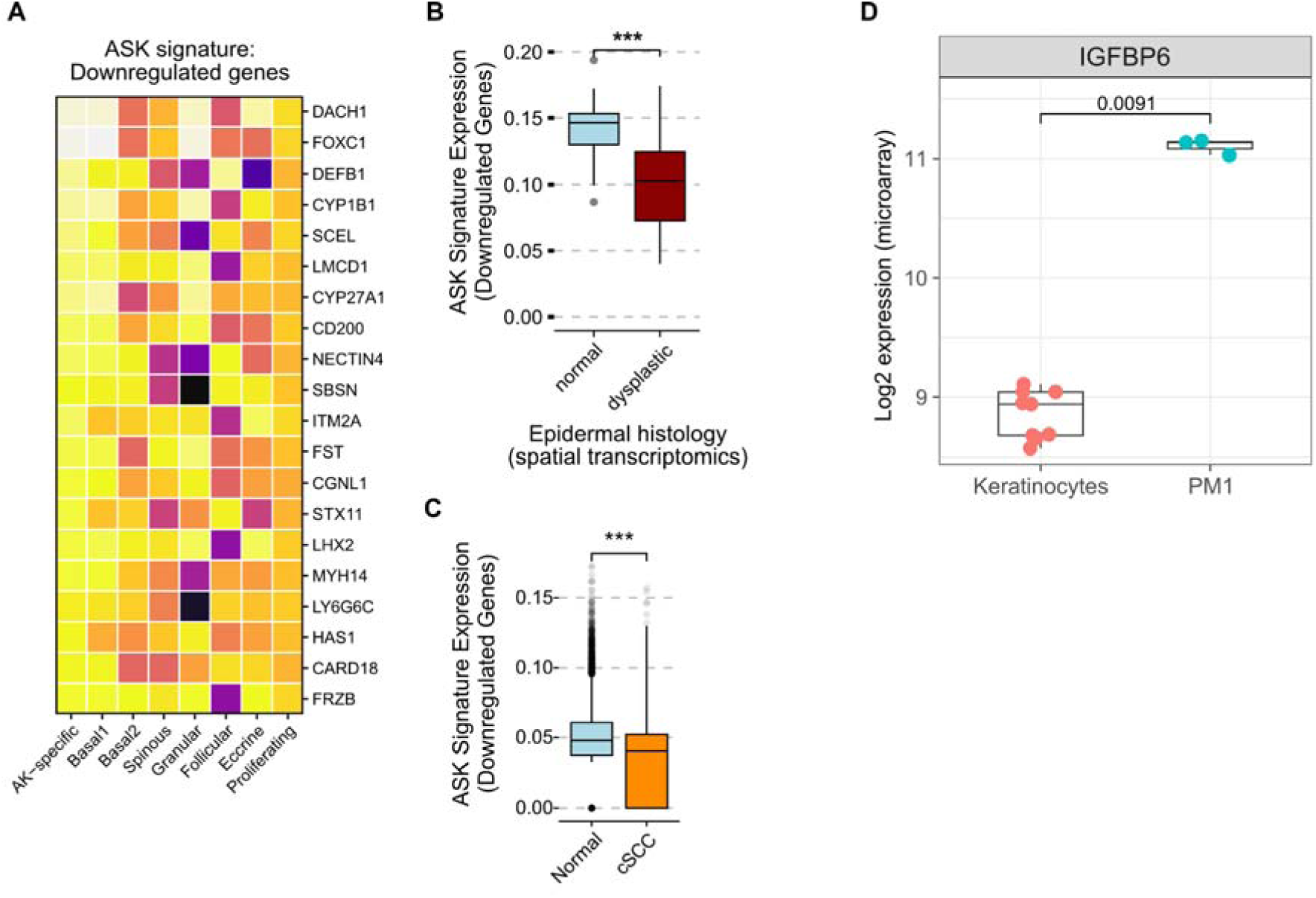
ASK signature downregulated genes across independent datasets. **(A)** Heatmap showing average expression of the top downregulated ASK signature genes across keratinocyte subclusters. **(B)** ASK signature enrichment scores (downregulated genes) in spatial transcriptomics spots annotated as morphologically normal or dysplastic epidermis (Wilcoxon rank-sum test). **(C)** ASK signature enrichment scores (downregulated genes) in normal and cSCC keratinocytes from Ji et al. 2020 (A. L. Ji et al., 2020) (Wilcoxon rank-sum test). **(D)** *IGFBP6* expression in normal keratinocytes and pre-malignant PM1 cells from published microarray dataset GSE98767 (Inman et al., 2018) (Wilcoxon rank-sum test).

**Supplementary Figure 5.**
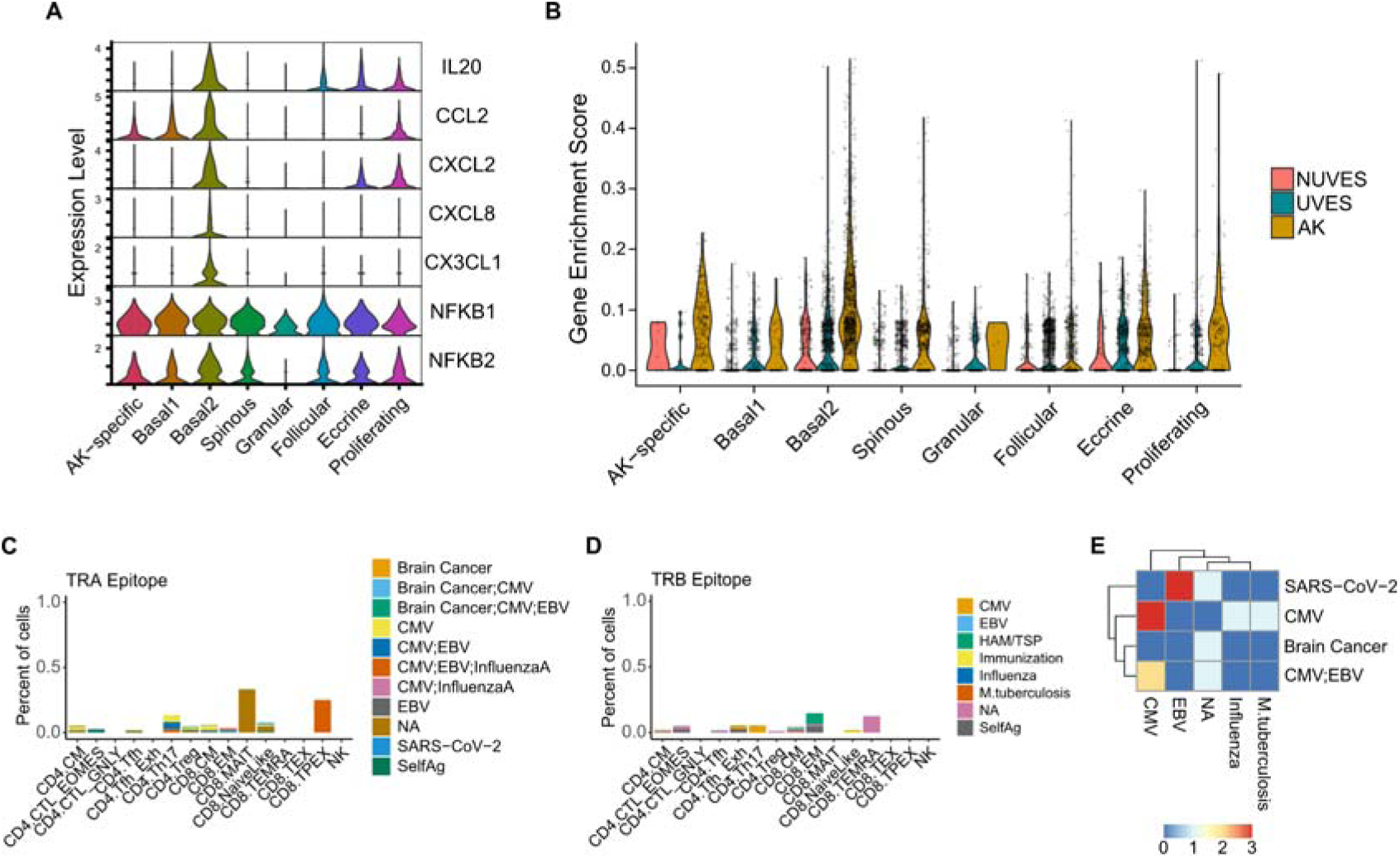
**(A)** Expression of inflammation-associated genes significantly overexpressed in basal2 keratinocytes. **(B)** Enrichment scores of the AK-associated keratinocyte signature from Li et al., 2023 (J. Li et al., 2023) across keratinocyte subclusters, by condition. **(C)** Predicted epitopes recognized by the T cell receptor (TCR) alpha chain (TRA) by T cell subtype. **(D)** Predicted epitopes recognized by the TCR beta chain (TRB) by T cell subtype. **(E)** Correlation between TRA- and TRB-predicted epitopes within individual TCRs.

